# Soluble P2X7 receptor is elevated in the plasma of COVID-19 patients and correlates with disease severity

**DOI:** 10.1101/2022.03.04.483019

**Authors:** Julio García-Villalba, Laura Hurtado-Navarro, Alejandro Peñín-Franch, Cristina Molina-López, Laura Martínez-Alarcón, Diego Angosto-Bazarra, Alberto Baroja-Mazo, Pablo Pelegrín

**Affiliations:** Biomedical Research Institute of Murcia (IMIB-Arrixaca), University Clinical Hospital Virgen Arrixaca, 30120 Murcia, Spain; Department of Biochemistry and Molecular Biology B and Immunology, Faculty of Medicine, University of Murcia, 30120 Murcia, Spain

**Keywords:** COVID-19, SARS-CoV-2, inflammation, P2X7, purinergic receptors, pyroptosis

## Abstract

Inflammation is a tightly coordinated response against bacterial and viral infections, triggered by the production of pro-inflammatory cytokines. SARS-CoV-2 infection induces COVID-19 disease, characterized by an inflammatory response mediated through the activation of the NLRP3 inflammasome, which results in the production of IL-1β and IL-18 along with pyroptotic cell death. The NLRP3 inflammasome could be also triggered by sterile danger signals such as extracellular ATP triggering the purinergic P2X7 receptor. Severe inflammation in the lungs of SARS-CoV-2 infected individuals is associated with pneumonia, hypoxia and acute respiratory distress syndrome, these being the causes of death associated with COVID-19. Both the P2X7 receptor and NLRP3 have been considered as potential pharmacological targets for treating inflammation in COVID-19. However, there is no experimental evidence of the involvement of the P2X7 receptor during COVID-19 disease. In the present study we determined the concentration of different cytokines and the P2X7 receptor in the plasma of COVID-19 patients and found that along with the increase in IL-6, IL-18 and the IL-1 receptor antagonist in the plasma of COVID-19 patients, there was also an increase in the purinergic P2X7 receptor. Increase in COVID-19 severity and C-reactive protein concentration positively correlated with increased concentration of the P2X7 receptor in the plasma, but not with IL-18 cytokine. The P2X7 receptor was found in the supernatant of human peripheral blood mononuclear cells after inflammasome activation. Therefore, our data suggest that determining levels of the P2X7 receptor in the plasma could be a novel biomarker of COVID-19 severity.

## Introduction

Coronavirus disease 2019 (COVID-19) develops after infection with the severe acute respiratory syndrome coronavirus 2 (SARS-CoV-2) and, according to the Johns Hopkins University COVID-19 dashboard, as of 10 February 2022 there have been over 400 million infections and 5.7 million deaths worldwide (coronavirus.jhu.edu/map.html). SARS-CoV-2 primarily infects cells in the respiratory tract and causes different degrees of symptomatology, with the severe cases developing into pneumonia with hypoxia and acute respiratory distress syndrome (ARDS) [1]. The lungs of COVID-19 patients with ARDS show considerable infiltration of neutrophils and monocytes and an increase in the concentration of several pro-inflammatory cytokines in the blood [1–6]. The nucleotide-binding oligomerization domain, leucine rich repeat and pyrin domain containing-protein 3 (NLRP3) inflammasome controls the release of bioactive interleukin (IL)-1β and IL-18 cytokines by activating caspase-1 and inducing pyroptosis cell death [7,8]. Caspase-1 also cleaves gasdermin D (GSDMD) that induces pores in the plasma membrane allowing the release of IL-1β and IL-18 [7,9,10]. The NLRP3 inflammasome is activated by different signals, including triggering of the P2X7 receptor by extracellular ATP, the phagocytosis of particulate matter and infection with specific viruses or bacteria [11–13]. The SARS-CoV-2 virus activates NLRP3 [14–18], and this inflammasome has been implicated in the production of pro-inflammatory cytokines in severe cases of COVID-19 and is thus directly associated with the severity of ARDS [15,19,20]. A clinical trial in COVID-19 patients with anakinra (the recombinant IL-1 receptor antagonist) has found that it decreases the severity of the respiratory failure by reducing the inflammatory response [21]. Mechanistically, the SARS-CoV-2 viroporin encoded by *ORF3a* triggers cellular K^+^ efflux and activates NLRP3 [22], with decreased intracellular K^+^ being a well-known activator of the NLRP3 inflammasome [23]. Also, the SARS-CoV-2 N protein and the *ORF8* encoded protein are able to interact with NLRP3 and induce its activation [24,25]. On one hand, during SARS-CoV-2 infection there is a reprogramming of the host macrophages, which means that they can be primed for the NLRP3 inflammasome by the virus spike (S) protein [26]. On the other hand, the SARS-CoV-2 virus blocks the inflammasome pathway at different levels; for example, the envelope (E) protein impairs NLRP3 priming [27], the non-structural proteins 1 and 13 suppress the activation of the NLRP3 inflammasome [28] and the N protein cleaves GSDMD within the N-terminal lytic domain, thus inactivating pyroptosis [29]. Therefore, there is a fine balance between positive and negative routes during NLRP3 inflammasome activation in cases of SARS-CoV-2 infection. Different review studies also suggest that the P2X7 receptor could be an important molecular pathway activated during COVID-19 disease, partly due to the activation of the NLRP3 inflammasome [30,31]. Activation of P2X7 also leads to the cellular shedding of this receptor [32] and, during different inflammatory diseases, an increase in the concentration of a soluble form of the receptor in the plasma [32–34]. However, in SARS-CoV-2 infection there is no experimental evidence regarding the potential involvement of P2X7 receptor.

In the present study, we have found that during active COVID-19 disease there are elevated concentrations in the plasma of IL-18, the IL-1 receptor antagonist (IL-1Ra), and a soluble form of the purinergic P2X7 receptor. IL-1Ra and the P2X7 receptor both increased with the severity of COVID-19 symptoms, but only the P2X7 receptor correlated with CRP concentration in the plasma of COVID-19 patients.

## Results

Plasma samples were collected from a cohort of 208 individuals with confirmed SARS-CoV-2 infection and different degrees of COVID-19 disease gravity between September 2020 and March 2021 in the Region of Murcia in the southeast of Spain (**Table 1**). The largest group among these individuals were those with moderate COVID-19 symptoms (N= 128), followed by those with mild symptoms (N= 41) and those with severe symptoms (N= 39). The control cohort used samples that had been collected during the period 2016-2019 (pre-COVID-19 pandemic) also in the Region of Murcia in the southeast of Spain. These samples were gathered from 69 individuals without SARS-CoV-2 infection and without inflammatory or infection symptoms (**Table 1**). The level of the acute phase C-reactive protein (CRP) was above the standard threshold in the plasma of 90.9% of the patients infected with SARS-CoV-2 (**Fig. 1A**). However, procalcitonin (PCT) was above the standard healthy threshold in only 10.6% of the COVID-19 patients (**Fig. 1A**), whereas ferritin was above the standard healthy threshold in 71.2% of the COVID-19 patients (taking into account male and female thresholds, **Fig. 1A**). As expected, IL-6 cytokine was also elevated in the plasma of patients infected with SARS-CoV-2 when compared to healthy control individuals (**Fig. 1B**); and IL-15, IL-1RA and IL-18 were also elevated (**Fig. 1B**), a finding that is in line with previous publications [3,16]. However, other cytokines or chemokines, such as CCL2/MCP-1 and IL-2, were not elevated in the plasma of the cohort of SARS-CoV-2 infected patients when compared to healthy individuals (**Fig. 1C**). Increases in these have been associated with mortality only in COVID-19 patients [3], and our cohort confirms that CCL2/MCP-1 was also significantly increased in the COVID-19 patients who die (237.4±12.4 *vs*. 421.9±59.0 pg/ml CCL2 mean±sem in COVID-19 survival *vs*. exitus; *p*= 0.018 Mann-Whitney test). However, values for IL-2 failed to associate with COVID-19 mortality (not shown), and this could probably due because IL-2 concentration was very low, close to detection threshold.

**Table 1.**
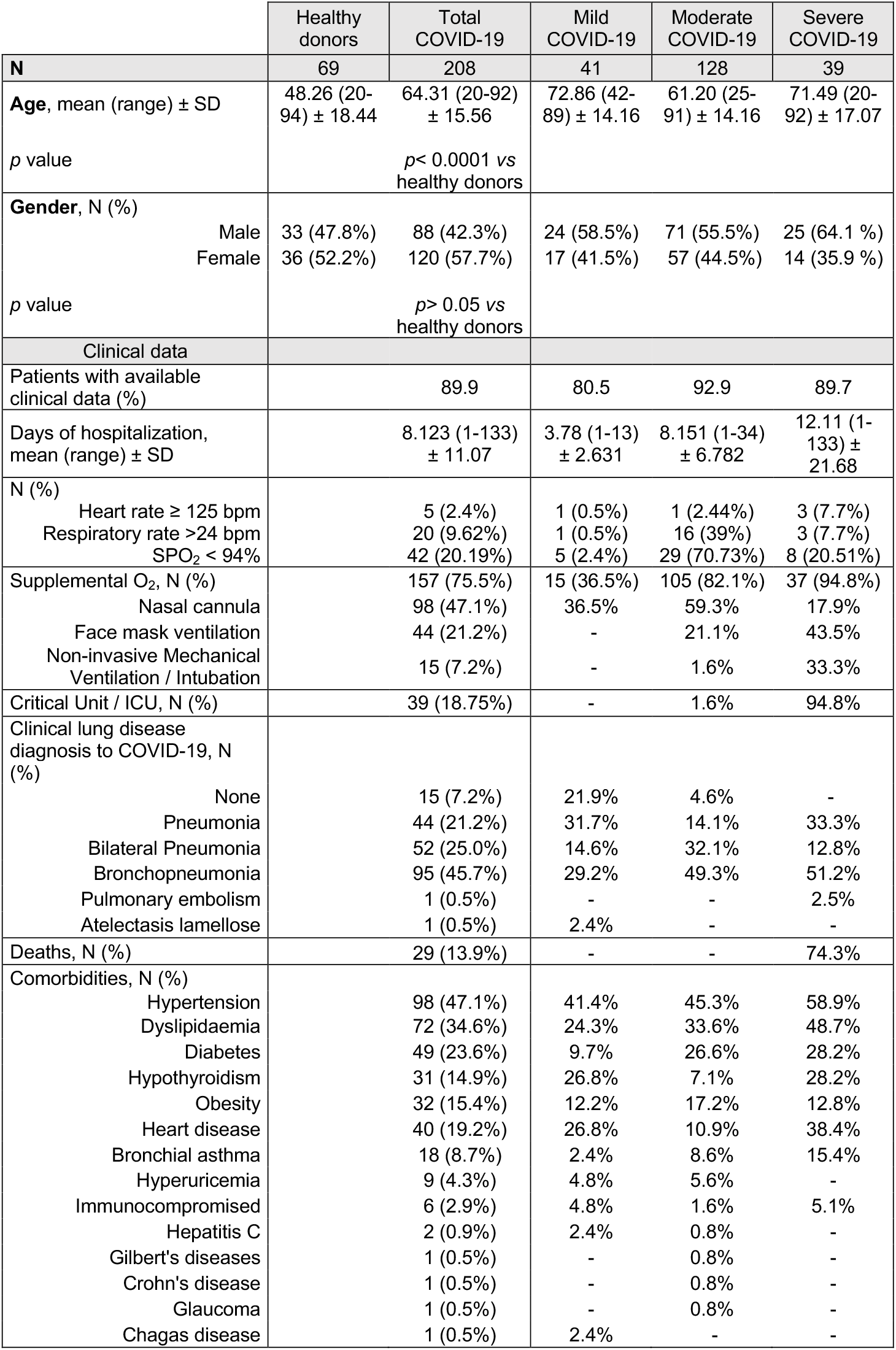
Demographic and clinical characteristics of the individuals included in this study.

**Figure 1.**
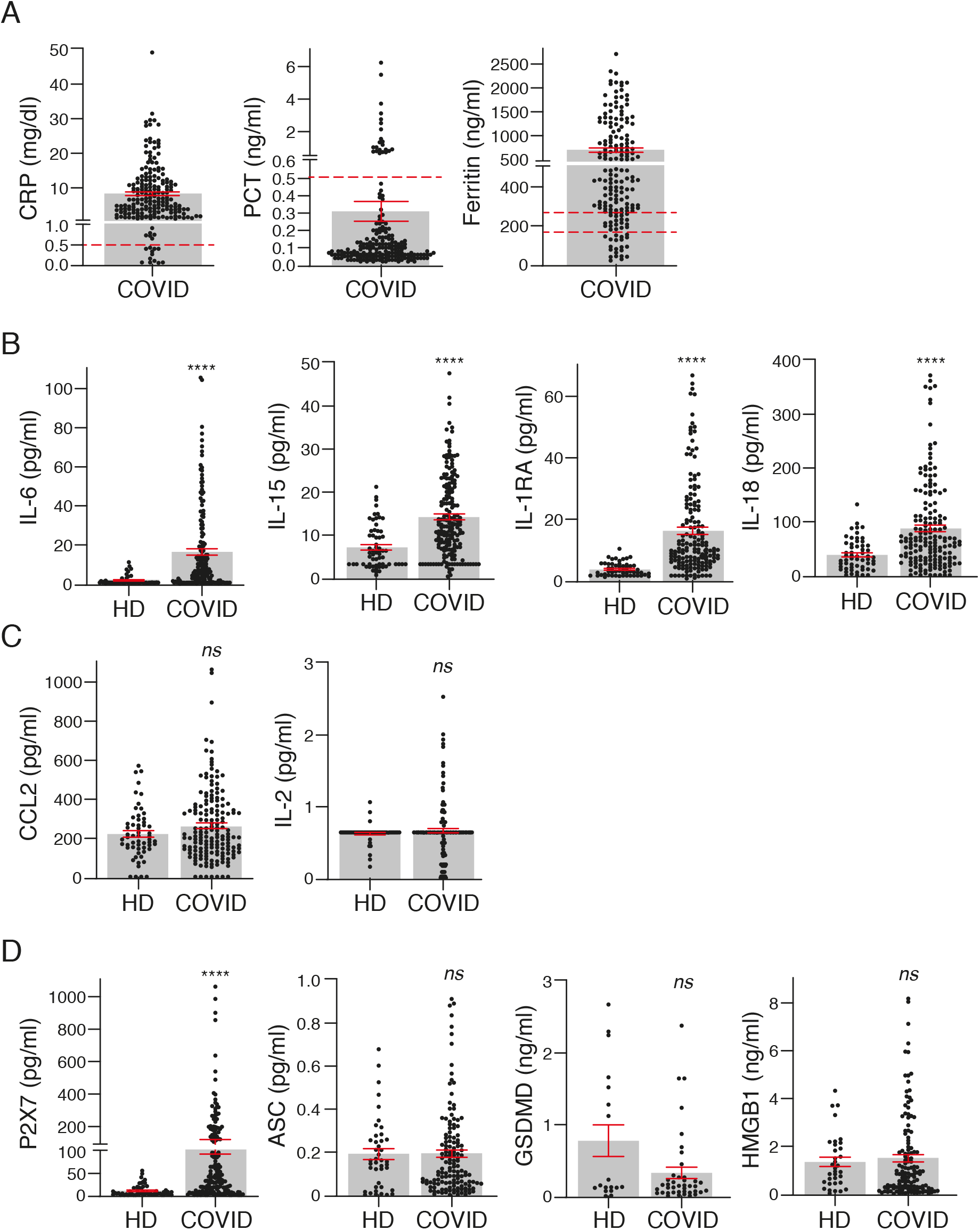
Inflammatory cytokines and biomarkers are markedly induced in COVID-19 patients. (**A**) Concentration of C-reactive protein (CRP), procalcitonin (PCT) and ferritin in plasma of COVID-19 patients. (**B-C**) Plasma concentration of IL-6, IL-15, IL-1RA, IL18 (**B**), CCL2, and IL-2 (**C**) in healthy donors (HD) and COVID-19 patients. (**D**) Plasma concentration of P2X7 receptor and markers of pyroptosis (ASC, GSDMD, and HMGB1) in healthy donors (HD) and COVID-19 patients. Each dot represents an individual patient; data are represented as mean (grey bars) ± s.e.m; dotted lines represent the normal range for each parameter analysed (upper line in Ferritin’s graph correspond to normal range for healthy men and bottom line for healthy woman); For (**B-D**) \*\*\*\**p* < 0.0001; *ns*, no significant difference (p > 0.05) with Mann-Whitney test.

The presence of the soluble P2X7 receptor was significantly increased in the plasma of SARS-CoV-2 patients when compared to healthy individuals (**Fig. 1E**). However, other markers of pyroptosis (HMGB1, ASC and GSDMD) were not elevated in the plasma of SARS-CoV-2 infected patients when compared to healthy individuals (**Fig. 1E**). Of particular note is that only 27.5% of healthy individuals and 25% of COVID-19 patients presented a positive detection of GSDMD in the plasma, suggesting that GSDMD was under the detection threshold in the majority of individuals and irrespective of the infection. Similarly, HMGB1 was detected in only 58% of healthy donors and in 74% of COVID-19 patients. Detection rates of all the other identified proteins were very similar among healthy donors and COVID-19 patients, with a detection range above 80% of the samples analysed.

The concentration of CRP in the plasma of COVID-19 patients significantly increased in line with the categories of illness severity (**Fig. 2A**). Illness severity categories were defined as mild, moderate and severe/critical in line with the clinical spectrum of SARS-CoV-2 infection defined in the latest COVID-19 treatment guidelines published by the National Institute of Health. A total of 41 mild, 128 moderate and 39 severe COVID-19 cases were identified (**Table 1**). However, PCT was significantly elevated in only the severe COVID-19 group (**Fig. 2A**) and ferritin was increased in the mild and severe groups (**Fig. 2A**).

**Figure 2.**
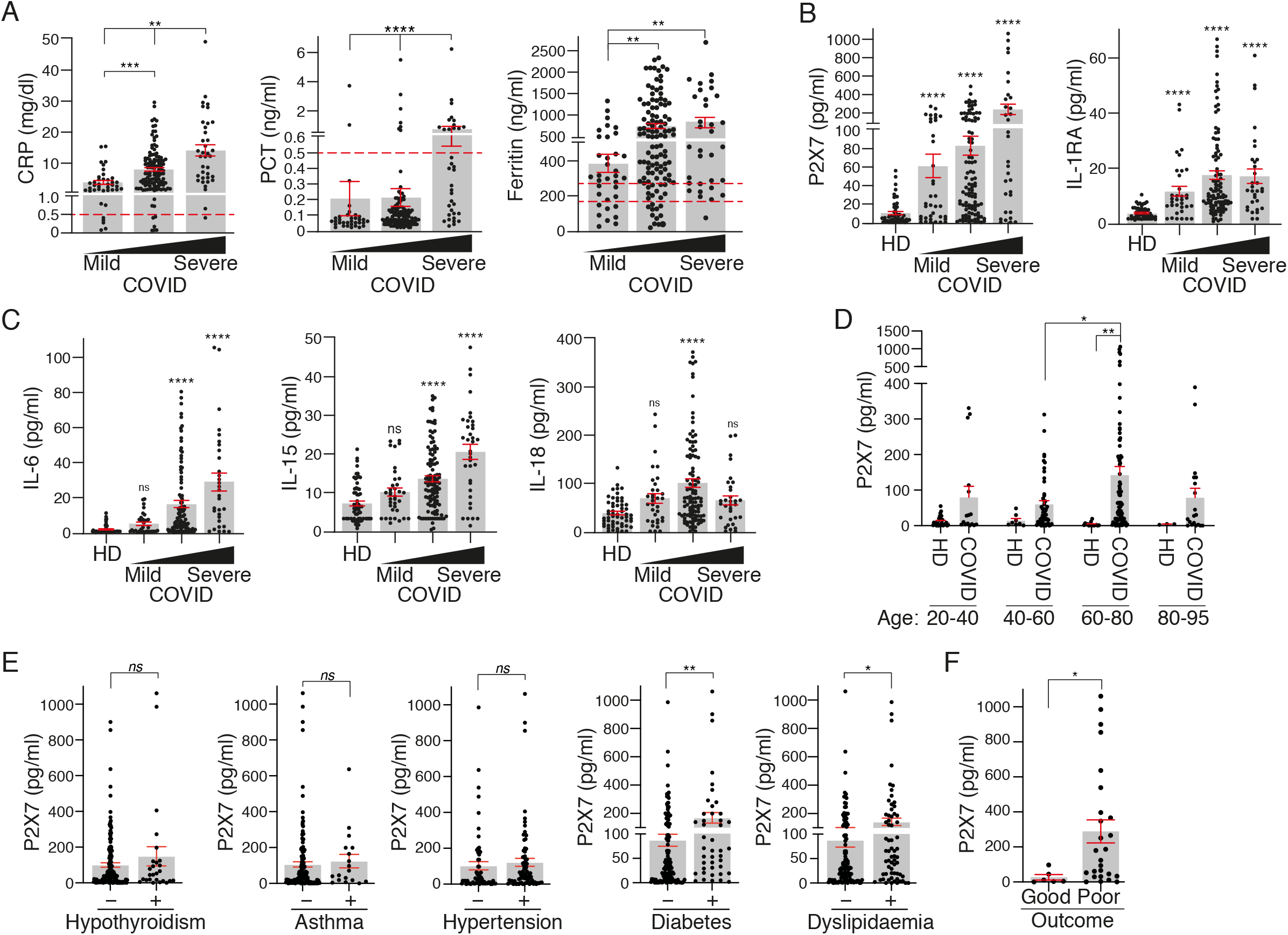
P2X7 receptor and IL-1RA increase with COVID-19 disease severity. (**A**) Plasma concentration of C-reactive protein (CRP), procalcitonin (PCT) and ferritin in association with the severity of COVID-19 manifestation (mild, moderate or severe). (**B,C**) Concentration of P2X7 receptor, IL-1RA (**B**), IL-6, IL-15 and IL-18 (**C**) in plasma of healthy donors (HD) and COVID-19 patients in association with the severity of disease manifestation (mild, moderate or severe). **(D)** Concentration of P2X7 receptor in the plasma of HD and COVID-19 patients in different age groups. (**E**) Concentration of P2X7 receptor in the plasma of COVID-19 patients with hypothyroidism, asthma, hypertension, diabetes or dyslipidaemia. (**D**) Concentration of P2X7 receptor in the plasma of severe COVID-19 patients with good or poor outcome (death or prolonged intensive care unit stay). Each dot represents an individual patient; data are represented as mean (grey bars) ± s.e.m; dotted lines represent the normal range for each parameter analysed (upper line in the Ferritin graph corresponds to normal range for healthy men and the bottom line corresponds to normal range for healthy woman); \**p* < 0.05; \*\**p* < 0.005; \*\*\**p* < 0.0005; \*\*\*\**p* < 0.0001; *ns*, no significant difference (*p* > 0.05) with Kruskal-Wallis test for A-D and Mann-Whitney test for E,D.

The P2X7 receptor and IL-1RA were significantly increased in the plasma of COVID-19 patients across all severity groups when compared to healthy controls (**Fig. 2B**). IL-6 and IL-15 were not increased in the mild COVID-19 group, but were significantly elevated in the moderate and severe groups (**Fig. 2C**). IL-18 was significantly increased only in moderate COVID-19 patients (**Fig. 2C**). The increase in the P2X7 receptor in the plasma of COVID-19 patients was higher in the 60 to 80 year-old group (**Fig. 2D**) and in COVID-19 patients with diabetes or dyslipidaemia (**Fig. 2E**). However, P2X7 receptor concentration in the plasma was not affected in COVID-19 patients with hypothyroidism, asthma or hypertension (**Fig. 2E**). Furthermore, P2X7 receptor was significantly increased in the plasma of COVID-19 severe patients who had a poor outcome (death or prolonged intensive care unit stay) (**Fig. 2F**) and a multivariate logistic regression analysis showed that an increase of the concentration of soluble P2X7 is associated with severe COVID-19 symptoms (OR = 1.04; [95% CI: 1.013 – 1.060]; *p*= 0.002). This multivariate analysis also revealed that patients affected by asthma increases 4.4 times of having severe COVID-19 (OR = 4.4; [95% CI: 1.575 – 12.186]; *p*= 0.005) or age increases 4.2 times of having severe COVID-19 (OR = 4.2; [95% CI: 1.597 – 11.139]; *p=* 0.004).

The amount of soluble P2X7 receptor and the concentration of the cytokines IL-6 and IL-15 positively correlated with the amount of CRP in the plasma (**Fig. 3A,B**). The positive correlation between the P2X7 receptor and CRP increased in the moderate and severe COVID-19 groups, but in contrast, the positive IL-6 and IL-15 correlation with CRP decreased in these groups (**Fig. 3C**). Neither IL-1RA nor IL-18 cytokines correlated with CRP when all COVID-19 patients were analysed (**Fig. 3A,B**) or when the moderate and severe groups were separated (not shown). Although CRP also positively correlated with ferritin concentration (**Fig. 3D**), ferritin had a weaker correlation with the P2X7 receptor and no correlation with IL-18 (**Fig. 3D**). However, the soluble P2X7 receptor positively correlated with the different cytokines analysed, the best correlation being when the presence of IL-18 increased in the plasma (**Fig. 3E**). This suggests that soluble P2X7 receptor increases with the severity of COVID-19 illness.

**Figure 3.**
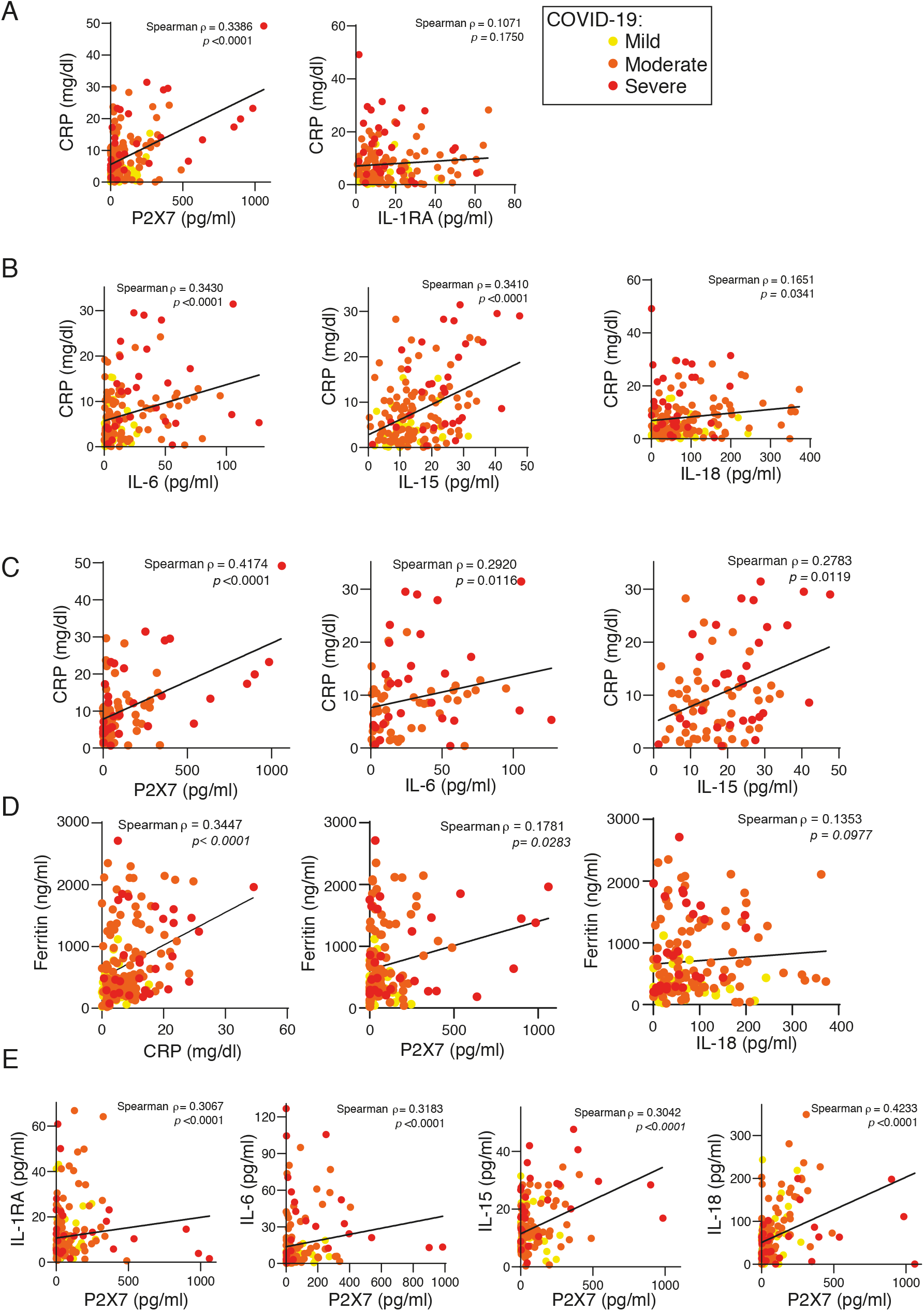
P2X7 receptor detected in the plasma of COVID-19 patients correlates with C-reactive protein. (**A,B**) Correlation between the concentration of C-reactive protein (CRP) and the quantification of the P2X7 receptor, IL-1RA (**A**), IL-6, IL-15 and IL-18 (**B**) in plasma from COVID-19 patients; disease severity is represented with yellow circles (mild), orange circles (moderate) and red circles (severe). (**C**) Correlation between the concentration of CRP and the quantification of the P2X7 receptor, IL-6 and IL-15 in plasma from COVID-19 patients with moderate (orange circles) and severe (red circles) COVID-19 manifestation. (**D**) Correlation between the concentration of ferritin and CRP, P2X7 receptor, and IL-18 in plasma from COVID-19 patients; disease severity is represented with yellow circles (mild), orange circles (moderate) and red circles (severe). (**E**) Correlation between the concentration of the P2X7 receptor and IL-1RA, IL-6, IL-15 and IL-18 from plasma of COVID-19 patients; disease severity is represented with yellow circles (mild), orange circles (moderate) and red circles (severe). Each dot represents an individual patient; Spearman correlation for all panels.

P2X7 receptor release has been observed in human macrophages after treatment with the semi-selective P2X7 receptor agonist benzoyl ATP [32]; we therefore asked if ATP treatment of human peripheral blood mononuclear cells (PBMCs) would also result in the shedding of the P2X7 receptor. Our results showed that ATP was able to increase the concentration of the P2X7 receptor found in the supernatant of LPS-treated PBMCs (**Fig. 4**). This LPS and ATP treatment also induces the activation of the NLRP3 inflammasome [11], as was evidenced by the release of IL-1β (**Fig. 4**). Furthermore, the P2X7 receptor and IL-1β were also released after the activation of a P2X7 receptor independent inflammasome, such as the Pyrin inflammasome (**Fig. 4**). The Pyrin inflammasome is activated by toxins that block RhoA GTPase, such as *Clostridium difficile* enterotoxin B (TcdB). RhoA blockage leads to the dephosphorylation of Pyrin and liberates it from the inhibitory 14-3-3 protein, which in turn leads to the activation of the Pyrin inflammasome [35]. These results support the notion that inflammasome-dependent caspase-1 activation is linked to the shedding of the P2X7 receptor from human blood cells, and given the inflammasome is activated during COVID-19 infection [14–16,18,20,26], this could be the cause of the elevated levels of P2X7 receptor in the plasma of these patients.

**Figure 4.**
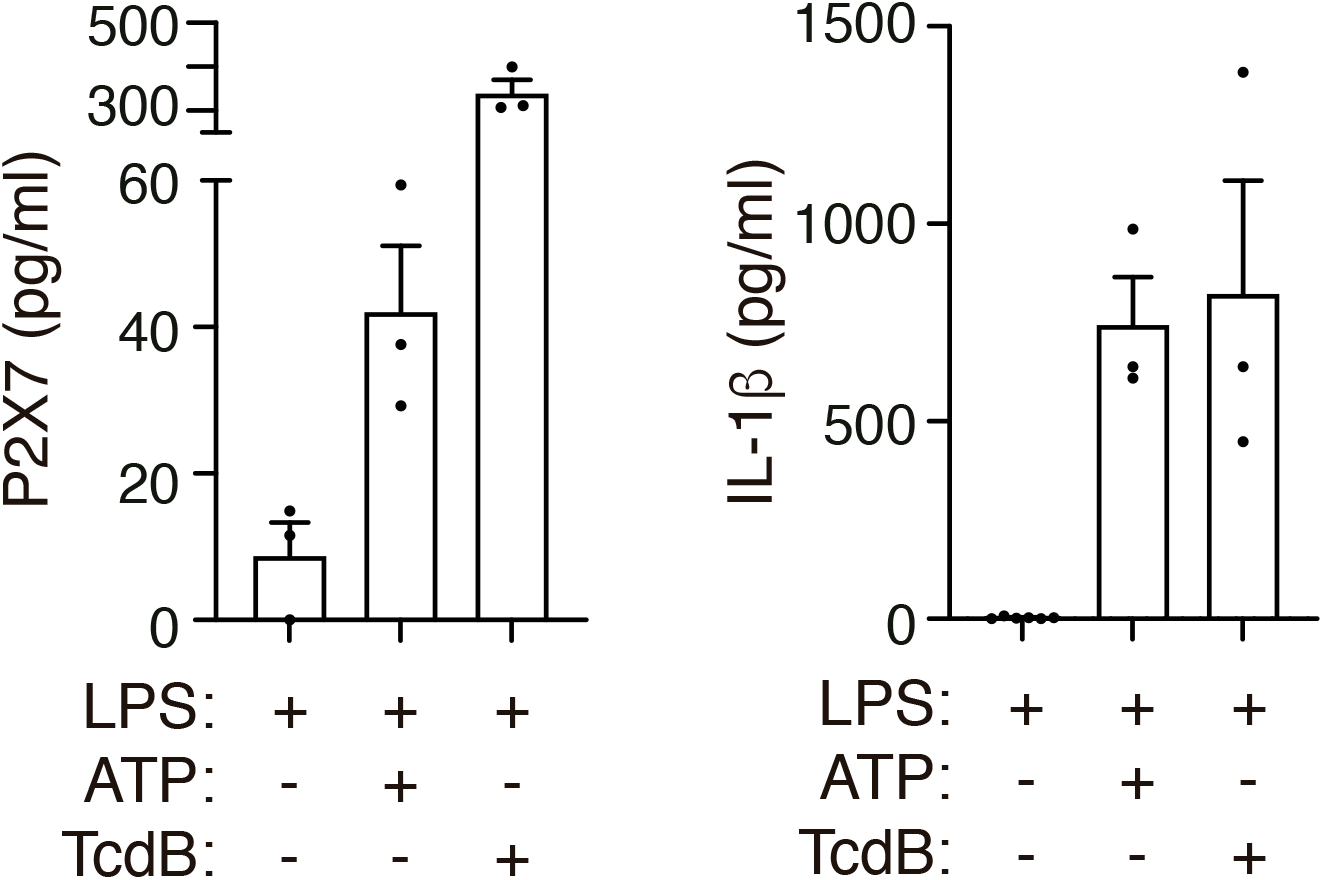
Treatment with ATP or *Clostridium difficile* toxin B induces the release of P2X7 receptor. P2X7 receptor (left) or IL-1β (right) from human peripheral blood mononuclear cells from healthy individuals treated for 3 h with LPS (1.6 µg/ml) and then treated with ATP (5 mM) for 45 min or with *Clostridium difficile* toxin B (TcdB, 1 µg/ml) for 1 h. Each dot represents data from an independent individual.

## Discussion

Severe COVID-19 symptoms are related to an exacerbated inflammatory response mediated by the production of several pro-inflammatory cytokines [1,3]. In the present study, we corroborate in a cohort of COVID-19 patients from the southeast of Spain an elevated concentration of the cytokines IL-6, IL-15, IL-18 and IL-1RA. Together with these cytokines, we found that levels of the purinergic receptor P2X7 also increase in the plasma of symptomatic COVID-19 patients, that these levels increase as the severity of the disease increases and that they correlate with the levels of the acute phase protein CRP, different cytokines and poor outcome. However, our cohort present a relatively low morality and association of the concentration of P2X7 receptor in the plasma with mortality would require further validation in larger cohorts. The activation of the P2X7 receptor leads to the activation of the NLRP3 inflammasome, the release of the IL-1β and IL-18 cytokines [11,13], and the shedding of the P2X7 receptor in microparticles [32]. Soluble P2X7 receptor found in cell supernatants is therefore associated with extracellular vesicles [32], and is therefore expected to be similar to the cellular plasma-membrane bound receptor. However, there is no data on the potential function of this shed form of the P2X7 receptor in extracellular vesicles, although circulatory exosomes from COVID-19 patients are able to activate the NLRP3 inflammasome [36]. In the present study, we show that human peripheral blood mononuclear cells release the P2X7 receptor when treated with ATP. This suggests that potential NLRP3 inflammasome activation and subsequent pyroptosis could be partially responsible for the release of microparticles loaded with P2X7 receptor, because the endosomal sorting complexes required for transport (ESCRT) machinery induces the shedding of GSDMD-damaged membranes [7,37]. In support of this hypothesis, we also describe in this study how the activation of the Pyrin inflammasome by TcdB and the subsequent pyroptosis (which is independent of the P2X7 receptor) also leads to the release of the P2X7 receptor. Increased concentrations of soluble P2X7 receptor in the blood have been described as positively correlating with the concentration of CRP [32,34], with the soluble P2X7 receptor being elevated during infection and sepsis [32,33], and during temporal lobe epilepsy [34]. Our results agree with those previous studies, and describe how infection with SARS-CoV-2 also increases the soluble P2X7 receptor in the plasma, which positively correlates with CRP concentration. In fact, recent studies have reported the activation of the inflammasome during SARS-CoV-2 infection and relate it to the severity of the COVID-19 infection [18]. All this evidence suggests that detecting the P2X7 receptor could be a promising novel blood-biomarker for inflammatory processes and help diagnose and determine the severity of COVID-19. The increase in the P2X7 receptor in the plasma of COVID-19 patients was higher in those with diabetes and dyslipidaemia, which in turn associates the P2X7 receptor with metabolic diseases [38,39]. In addition, targeting the P2X7 receptor with drug-like pharmacological antagonists has been suggested as a promising strategy for treating severe-inflammatory and neurological complications in COVID-19 patients [30,40].

The concentration of IL-6 has been clinically used to evaluate the severity of COVID-19 [3]. Also, IL-6 has been used as a therapeutic target, and IL-6 blocking antibodies have been widely used to treat COVID-19 patients [41]. IL-6 is a cytokine produced downstream from IL-1β, whereas IL-1RA is a cytokine that blocks IL-1 signalling as it binds to the type I IL-1 receptor and prevents IL-1 from triggering its receptor [42]. In diseases satisfactorily treated with anti-IL-1 therapy, such as sepsis or autoinflammatory syndromes, the blood concentrations of IL-1β are very low and usually not detected [33,43]. According to previous publications and despite the evidence of inflammasome activation during COVID-19, IL-1β has not been detected in the blood samples of COVID-19 patients [5,16]. In this regard, detecting IL-1RA has been proposed as a systemic indicator of IL-1β production, and our study, in agreement with previous studyies [4,16], shows an important increase in IL-1RA in the plasma of COVID-19 patients. These results suggest that there is potential production of IL-1β during COVID-19 infection [6]. While the axes IL-1β/IL-6 link to CRP production in bacterial infections, IL-18 production links to transferrin in viral infection [44]. However, in the plasma of SARS-CoV-2 infected patients we found increased levels of both axes, IL-6/CRP and IL-18/ferritin; however, although IL-6 positively correlated with CRP, IL-18 failed to correlated with ferritin in COVID-19 patients. P2X7 correlated with CRP, a finding supported by previous studies [32,34], but not with ferritin, denoting that the shedding of the P2X7 receptor is somehow related to the IL-6/CRP inflammatory programme in both bacterial [45] and SARS-CoV-2 viral (this study) infections.

In conclusion, our study shows that the soluble P2X7 receptor concentration increases in the plasma of COVID-19 and positively correlates with disease severity and CRP protein concentration. This correlation was not found for IL-18 or other cytokines, although these were found to increase in the plasma of COVID-19 patients. Furthermore, the P2X7 receptor was released from human PBMCs upon inflammasome activation, suggesting that the P2X7 receptor could be a novel blood biomarker for COVID-19 severity with poor outcome.

## Material and methods

### Human samples

Samples and data from patients included in this study, who gave written informed consent, were collected, processed and provided by the *Biobanco en Red de la Región de Murcia*, BIOBANC-MUR, were registered on the *Registro Nacional de Biobancos* with registration number B.0000859, and were processed following standard operating procedures with the appropriate approval of the Ethical Committees (2021-5-10-HCUVA). The study included 208 recovered individuals who had previously given positive RT-PCR tests, positive serology, and/or positive rapid antigen tests for SARS-CoV-2. The plasma samples were collected in the days following hospital admission or diagnosis of SARS-CoV-2 infection. For research purposes across studies, patients with COVID-19 were divided according the clinical spectrum of SARS-CoV-2 infection into three categories of disease severity, namely mild, moderate and severe/critical, as described in the latest National Institute of Health COVID-19 treatment guidelines (www.covid19treatmentguidelines.nih.gov, accessed 11/25/2021). Mild illness is attributed to hospitalized patients who have any of the various symptoms associated to COVID-19 (such as headache, nausea, sore throat, fever, cough, loss of taste and smell, malaise or muscle pain) but who have no shortness of breath, dyspnea, oxygen supplementation or abnormal chest imaging. Moderate illness is attributed to hospitalized patients who show evidence of lower respiratory disease during clinical assessment or chest radiograph and who have an oxygen saturation (SpO_2_) ≥94% on room air at sea level requiring, in some cases, oxygen therapy. Severe/critical illness is attributed to hospitalized patients in the Critical Care Unit who have SpO_2_ <94% on room air at sea level, a respiratory rate > 24 breaths/min, highest level of supplemental oxygen required, bilateral infiltrates, and, individuals who die from respiratory failure caused by COVID-19 disease, septic shock and/or organ dysfunction. The degree of oxygen supplementation increases depending on the severity of the disease, although the oxygen needs vary depending on the degree of hypoxia in each patient. The negative control group consisted of 69 healthy blood donors from whom plasma samples were collected before the COVID-19 pandemic, meaning that they were free of COVID-19. Sera were stored at -80ºC until use. Mononuclear cells from healthy donors were purified after the donors had signed their informed consent agreement and whole peripheral blood samples were collected (*n=3*).

### Reagents

The different reagents used in this assay and their sources were: ultrapure *Escherichia coli* lipopolysaccharide (LPS) serotype 0111:B4 (InvivoGen), Adenosine 5’-triphosphate (ATP) from Sigma-Aldrich and *Clostridium difficile* toxin B (TcdB) from Enzo Life Sciences.

### PBMCs stimulation

Human peripheral blood mononuclear cells were collected using Ficoll Histopaque-1077 (Sigma-Aldrich) and cultured in Opti-MEM Reduced Serum Media (Invitrogen). PBMCs from patients were left unstimulated or stimulated with 1.6 μg/ml of LPS at 37 ºC and then subsequently stimulated with ATP 5 mM for 45 min or TcdB 1 μg/ml for 1h.

### ELISA

Individual culture cell-free supernatants were collected and clarified by centrifugation. The concentration of human soluble P2X7 was tested by ELISA following the manufacturer’s instructions (Cusabio). Plasma levels of human ASC, HMGB1, GSDMD and P2X7 were also tested by ELISA (Cusabio for ASC and P2X7, Aviva System Biology for GSDMD and Arigo Biolaboratories for HMGB1). Results were read in a Synergy Mx (BioTek) plate reader at 450 nm and corrected at 540 nm.

### Multiplex

Multiplexing in human serum for IL-15, IL-18, MCP-1, IL-2, IL-6 and IL-1RA was performed using the Luminex color-coded antibody-immobilized beads from Merck Millipore following the manufacture’s indications, and the results were analysed in a Luminex MAGPIX instrument (Luminex Corporation).

### Statistics and data analysis

Statistical analyses were performed using GraphPad Prism 9 (GraphPad Software Inc.). Normality of the samples was determined using the D’Agostino and Pearson omnibus K2 normality test. Outliers from data sets were identified by the ROUT method with *Q* = 1% and were eliminated from the analysis and representation. Non-parametric Mann-Whitney test was used to compare differences among two non-paired groups, and Kruskal-Wallis test was used to compare differences between three or more groups. The χ^2^-test was used to determine whether there was a significant difference between clinical variables among groups of patients. Non-parametric correlations were made according to the Spearman correlation coefficient. All data are shown as mean values and error bars represent standard error. To identify risk factors associated with SARS-CoV-2 severity multivariable analysis was done with SPSS software version 23 (SPSS Inc., Chicago, IL, USA). Age (groups of 20-60 and 60-95 years old), hypothyroidism, asthma, hypertension, diabetes and dyslipidaemia were reported as categorical variables and the concentration of soluble P2X7 receptor in the plasma was reported as continuous variable. We performed a multivariable logistic regression with those variables which significant difference compared using the χ^2^-test, being the dependent variable the SARS-CoV-2 severity (mild and moderate, vs severe). R^2^ Cox coefficient was used to assess the proportion of variation explained by the variables studied. For this context, a R^2^ Cox coefficient of 0 explain that there is no relationship between the variables and the variability in the data and, whereas a R^2^ Cox coefficient of 1 suggest that the variables in the regression explain 100 % of the variability in the data.

## Authors’ Contributions

JG-V, LH-N, AP-F and CM-L, experimental execution; JG-V, LH-N, and LM-A, sample collection; JG-V, LH-N, AP-F, CM-L, DA-B, AB-M and PP, data analysis; AB-M, DA-B and PP conceived the experiments; JG-V, LH-N and PP prepared the figures and wrote the paper; PP provided funding and overall supervision of the study.

## Acknowledgments

We are particularly grateful for the generous contribution of the donors and the collaboration of the Biobank Network of the Region of Murcia, BIOBANC-MUR, which is registered on the *Registro Nacional de Biobancos* with registration number B.0000859. BIOBANC-MUR is supported by the *Instituto de Salud Carlos III* (grant PT20/00109), by the *Instituto Murciano de Investigación Biosanitaria* and by the *Consejeria de Salud de la Comunidad Autónoma de la Región de Murcia*. We would like to thank all the members of Dr Pelegrin’s laboratory for comments and suggestions.

## References

1 Zhang, Q. et al. (2022) Human genetic and immunological determinants of critical COVID-19 pneumonia. Nature DOI: 10.1038/s41586-022-04447-0

2 Guo, Y.-R. et al. (2020) The origin, transmission and clinical therapies on coronavirus disease 2019 (COVID-19) outbreak – an update on the status. Mil. Med. Res. 7, 11

3 Abers, M.S. et al. (2021) An immune-based biomarker signature is associated with mortality in COVID-19 patients. JCI Insight 6,

4 Wilson, J.G. et al. (2020) Cytokine profile in plasma of severe COVID-19 does not differ from ARDS and sepsis. JCI insight 5, 1–6

5 Del Valle, D.M. et al. (2020) An inflammatory cytokine signature predicts COVID-19 severity and survival. Nat. Med. 26, 1636–1643

6 McElvaney, O.J. et al. (2020) Characterization of the Inflammatory Response to Severe COVID-19 Illness. Am. J. Respir. Crit. Care Med. 202, 812–821

7 Broz, P. et al. (2020) The gasdermins, a protein family executing cell death and inflammation. Nat. Rev. Immunol. 20, 143–157

8 Swanson, K. V. et al. (2019) The NLRP3 inflammasome: molecular activation and regulation to therapeutics. Nat. Rev. Immunol. 19, 477–489

9 Evavold, C.L. et al. (2018) The pore-forming protein gasdermin D regulates interleukin-1 secretion from living macrophages. Immunity 48, 35–44.e6

10 Xia, S. et al. (2021) Gasdermin D pore structure reveals preferential release of mature interleukin-1. Nature 593, 607–611

11 Di Virgilio, F. et al. (2017) The P2X7 Receptor in Infection and Inflammation. Immunity 47, 15–31

12 Christgen, S. et al. (2020) Toward targeting inflammasomes: insights into their regulation and activation. Cell Res. 30, 315–327

13 Pelegrin, P. (2021) P2X7 receptor and the NLRP3 inflammasome: Partners in crime. Biochem. Pharmacol. 187, 114385

14 Kroemer, A. et al. (2020) Inflammasome activation and pyroptosis in lymphopenic liver patients with COVID-19. J. Hepatol. 73, 1258–1262

15 Rodrigues, T.S. et al. (2021) Inflammasomes are activated in response to SARS-CoV-2 infection and are associated with COVID-19 severity in patients. J. Exp. Med. 218, e20201707

16 Junqueira, C. et al. (2021) SARS-CoV-2 infects blood monocytes to activate NLRP3 and AIM2 inflammasomes, pyroptosis and cytokine release. Res. Sq. DOI: 10.21203/rs.3.rs-153628/v1

17 Kucia, M. et al. (2021) An evidence that SARS-Cov-2/COVID-19 spike protein (SP) damages hematopoietic stem/progenitor cells in the mechanism of pyroptosis in Nlrp3 inflammasome-dependent manner. Leukemia DOI: 10.1038/s41375-021-01332-z

18 Vora, S.M. et al. (2021) Inflammasome activation at the crux of severe COVID-19. Nat. Rev. Immunol. 21, 694–703

19 Amin, S. et al. (2021) NLRP3 inflammasome activation in COVID-19: an interlink between risk factors and disease severity. Microbes Infect. 110, 104913

20 Courjon, J. et al. (2021) Heterogeneous NLRP3 inflammasome signature in circulating myeloid cells as a biomarker of COVID-19 severity. Blood Adv. 5, 1523–1534

21 Kyriazopoulou, E. et al. (2021) An open label trial of anakinra to prevent respiratory failure in covid-19. Elife 10, 1–21

22 Xu, H. et al. (2022) SARS-CoV-2 viroporin encoded by ORF3a triggers the NLRP3 inflammatory pathway. Virology 568, 13–22

23 Tapia-Abellán, A. et al. (2021) Sensing low intracellular potassium by NLRP3 results in a stable open structure that promotes inflammasome activation. Sci. Adv. 7, eabf4468

24 Wu, X. et al. (2021) Secreted ORF8 is a pathogenic cause of severe Covid-19 and potentially targetable with select NLRP3 inhibitors. bioRxiv DOI: 10.1101/2021.12.02.470978

25 Pan, P. et al. (2021) SARS-CoV-2 N protein promotes NLRP3 inflammasome activation to induce hyperinflammation. Nat. Commun. 12, 4664

26 Theobald, S.J. et al. (2021) Long-lived macrophage reprogramming drives spike protein-mediated inflammasome activation in COVID-19. EMBO Mol. Med. DOI: 10.15252/emmm.202114150

27 Yalcinkaya, M. et al. (2021) Modulation of the NLRP3 inflammasome by Sars-CoV-2 Envelope protein. Sci. Rep. 11, 24432

28 Kim, N.-E. et al. (2021) SARS-CoV-2 Nonstructural Proteins 1 and 13 Suppress Caspase-1 and the NLRP3 Inflammasome Activation. Microorganisms 9, 494

29 Ma, J. et al. (2021) SARS-CoV-2 nucleocapsid suppresses host pyroptosis by blocking Gasdermin D cleavage. EMBO J. 40, 1–17

30 Di Virgilio, F. et al. (2020) A rationale for targeting the P2X7 receptor in Coronavirus disease 19. Br. J. Pharmacol. 177, 4990–4994

31 Hasan, D. et al. (2021) A novel definition and treatment of hyperinflammation in COVID-19 based on purinergic signalling. Purinergic Signal. DOI: 10.1007/s11302-021-09814-6

32 Giuliani, A.L. et al. (2019) The P2X7 Receptor Is Shed Into Circulation: Correlation With C-Reactive Protein Levels. Front. Immunol. 10, 1–7

33 Martínez-García, J.J. et al. (2019) P2X7 receptor induces mitochondrial failure in monocytes and compromises NLRP3 inflammasome activation during sepsis. Nat. Commun. 10, 2711

34 Conte, G. et al. (2021) Circulating P2X7 Receptor Signaling Components as Diagnostic Biomarkers for Temporal Lobe Epilepsy. Cells 10, 2444

35 Masters, S.L. et al. (2016) Familial autoinflammation with neutrophilic dermatosis reveals a regulatory mechanism of pyrin activation. Sci. Transl. Med. 8, 332ra45

36 Sur, S. et al. (2022) Circulatory exosomes from COVID-19 patients trigger NLRP3 inflammasome in endothelial cells. bioRxiv DOI: 10.1101/2022.02.03.479081

37 Rühl, S. et al. (2018) ESCRT-dependent membrane repair negatively regulates pyroptosis downstream of GSDMD activation. Science 362, 956–960

38 Jain, S. and Jacobson, K.A. (2021) Purinergic signaling in diabetes and metabolism. Biochem. Pharmacol. 187, 114393

39 Solini, A. and Novak, I. (2019) Role of the P2X7 receptor in the pathogenesis of type 2 diabetes and its microvascular complications. Curr. Opin. Pharmacol. 47, 75–81

40 Ribeiro, D.E. et al. (2021) Hyperactivation of P2X7 receptors as a culprit of COVID-19 neuropathology. Mol. Psychiatry 26, 1044–1059

41 Cortegiani, A. et al. Rationale and evidence on the use of tocilizumab in COVID-19: a systematic review. Pulmonology 27, 52–66

42 Dinarello, C.A. (1997) Interleukin-1. Cytokine Growth Factor Rev. 8, 253–65

43 Baroja-Mazo, A. et al. (2014) The NLRP3 inflammasome is released as a particulate danger signal that amplifies the inflammatory response. Nat. Immunol. 15, 738–748

44 Slaats, J. et al. (2016) IL-1β/IL-6/CRP and IL-18/ferritin: Distinct Inflammatory Programs in Infections. PLoS Pathog. 12, 1–13

45 Holub, M. et al. (2013) Cytokines and chemokines as biomarkers of community-acquired bacterial infection. Mediators Inflamm. 2013, 190145

